# A pilot study to evaluate the effect of a novel calcium and vitamin D-containing oral bolus on serum calcium levels in Holstein dairy cows following parturition

**DOI:** 10.1101/520221

**Authors:** Daniel A. Shock, Steven M. Roche, Rachel Genore, Merle E. Olson

## Abstract

The initiation of lactation challenges the ability of the modern lactating cow to maintain calcium homeostasis, and typically results in a drop in blood calcium levels; leading to mobilization of calcium reserves from skeletal stores. As such, the recommendation to provide supplemental calcium at parturition to older cows has become an industry-standard practice. Mature cows were treated at calving and 12 hours later with either the novel calcium bolus (NB), or a commercially available calcium bolus (CB). Blood was collected from animals at 0, 1, 6, 12, 13, and 24 hours following calving, and the resulting serum samples were analyzed. Overall, there was no statistical difference between the NB and CB groups for blood calcium levels within the first 24 hours following parturition (P = 0.50). Cows in both groups experienced a significant increase in serum calcium by 1 hour after parturition; however, this increase was not sustained through subsequent sampling times. This pilot study demonstrates that both boluses have a similar effect in the elevation of blood calcium.

## Introduction

The initiation of milk production challenges the ability of the lactating dairy cow to maintain normal calcium concentrations. Once lactation begins, blood calcium levels drop leading to mobilization of calcium reserves from skeletal stores ^1,2^. Clinical and subclinical hypocalcaemia are common issues in the dairy industry ^3–5^, and both are associated with a variety of negative sequelae including an increased risk of displaced abomasum, mastitis, retained placenta, dystocia, and ketosis ^6,7^. Given these negative connotations, it is imperative to improve the calcium status of a newly freshened dairy cow.

Oral supplementation with calcium chloride and calcium sulfate in a fat coated bolus have been shown to have significant effects on improving calcium status in the period following calving ^8^. Clinically, Oetzel and Miller ^9^ found that the administration of the bolus to lame cows and high producing cows lead to fewer health events and higher milk production in those groups, respectively leading to an economic benefit of $3 to $8 per cow ^10^. There has been very strong adoption of the oral calcium supplement by dairy producers throughout North America; however, many of the alternative products to this product have not been tested to demonstrate similar efficacy in influencing blood calcium levels.

Therefore, the primary objective of this pilot study was to assess the ability of a novel oral calcium bolus containing calcium chloride, calcium sulfate, and vitamin D3 to influence the serum calcium level in recently fresh, mature Holsteins over the first 24 hours following parturition, relative to a positive control. We hypothesized that there would be no difference between the two oral boluses with respect to serum calcium levels over the first 24 hours following administration.

## Material and methods

### Ethics statement

The present field-based study was conducted in compliance with the research guidelines set forth by the Canadian Council on Animal Care after appropriate Animal Use Protocol review by the Institutional Animal Care and Use Committee (Alberta Agriculture, Airdrie Alberta, Canada).

### Sample size calculation

Based on previous research, oral calcium supplementation should increase the serum calcium concentration by 0.25 mmol/L (0.125 mmol/L standard deviation) relative to levels at calving ^11^. A total sample size of 20 mature cows - 10 each in the novel bolus (NB, 579.1 mg/g calcium chloride, 231.6 mg/g calcium sulfate, and 50,000 IU vitamin D3; Calboost, Solvet, Calgary, Alberta, Canada) and commercially available bolus (CB, containing 579.1 mg/g calcium chloride, 231.6 mg/g calcium sulfate; Bovikalc, Boehringer Ingelheim Vetmedica Inc., St. Joseph, MO, USA) groups was calculated, assuming a 95% confidence level and 80% power.

### Study herd and treatment protocols

The study was conducted at a commercial dairy farm milking 500 cows in southwestern Ontario, Canada, between March and September 2018. Average daily milk production per cow was 38 kg, at 3.9% fat and 3.1% protein. The herd averaged 155 days in milk. The dry cow ration contained straw, corn silage, and protein supplement, and is considered a low calcium, low phosphorous, low potassium, high magnesium diet that did not contain anionic salts. Historically, the farm averaged less than 5% incident risk of clinical hypocalcemia. This farm was chosen for proximity to the veterinary clinic that assisted with sample handling and analysis, as well as technical capability to collect and record relevant samples and data.

Cows were randomly allocated to receive either the NB or CB according to a randomization sheet that was generated using the RAND command in Microsoft Excel. In this experiment, a positive control was used, as the practice of oral calcium supplementation at calving is an industry-standard recommendation, and few producers capable of conducting field-based research are willing to accept the risk of not prophylactically treating all mature cows with oral calcium. First lactation cows, and those animals whose calvings were unobserved were excluded from enrollment. The farm staff was responsible for bolus administration, blood sample collection and storage, and reporting any disease events. They were blinded to the treatment groups throughout the trial.

The bolus was given orally at calving and again at 12 hours post-calving. Blood samples were collected via venipuncture of the coccygeal vein. Blood was taken at 0, 1, 6, 12, 13, and 24 hours relative to calving. Due to practicalities associated with sampling enrolled cows, the sampling window could vary by ± 1 hour of the protocol designation. Samples were allowed to clot, centrifuged for 5 minutes at 800 x g, and serum was transferred into sterile storage vials. The vials were stored at −20°C until submitted for analysis. Samples were analyzed to determine total serum calcium, phosphorus, and magnesium via a photometric method (Abaxis Vetscan vs2, Abaxis Inc., Union City, CA, USA).

Following treatments, participating farm personnel were instructed to record treatment assignment and all pertinent health and disease events (including death and culling) for all cows that had calved after study commencement.

### Data collection

Variables of interest for analysis included date and time of calving, calcium bolus administration, and blood collection, lactation number, dystocia, non-lactating days prior to calving, disease dates, culling/death dates, and occurrence of twin pregnancies. All results were transcribed into Excel (Microsoft Corporation, 2010, Redmond, WA, USA).

### Statistical analysis

All datasets were imported as comma-separated files into the Stata 14 (StataCorp LP, College Station, TX, USA) statistical software program for analysis. The cow was considered the unit of analysis for this study.

#### Descriptive statistics and univariable analysis

Descriptive statistics were generated for the final dataset. Serum calcium, phosphorous, and magnesium levels were explored through graphical plots. Specific comparisons between the serum calcium, days dry, lactation, season of sampling, and the presence of twins at calving in NB and CB cows were made, with univariable statistical comparisons made using Fisher exact tests or Kruskall-Wallis one-way analysis of variance.

Statistical comparisons were made between potential predictor variables and outcomes of interest. For this study, the outcome of interest was serum calcium level (in mmol/L). Univariable linear regression modelling was employed to study variables of interest. Variables explored for each model included: treatment group (NB vs CB), lactation group (2 and over 3), non-lactating days prior to parturition, season of calving, sampling time (0, 1, 6, 12, 13, and 24 hours relative to calving), dystocia (0 = eutotic calving, 1 = dystotic calving), and presence of twins at calving. Variables that had moderate statistical associations with the outcome of interest (defined at a liberal P-value < 0.2) were included in subsequent multivariable models.

#### Multivariable regression analysis

A repeated-measures mixed linear regression model was constructed, using random effects to model sample number within cow, using a first-order autoregressive correlation structure between time points.

A step-wise backwards elimination process was employed, where variables identified as potentially associated with the outcome of interest in the initial univariable screening were included in a full multivariable model. Those variables having P-values for partial F-tests or type III tests of fixed effects greater than 0.05 were eliminated from the model after assessing whether they were part of biologically plausible interaction terms or had a confounding effect on the outcome of interest (e.g. a > 20% change in coefficient values when the term is removed from the model). Continuous variables were assessed for linearity with predicted model outcomes through visually inspecting a LOWESS curve (local weight scatterplot smoothing) for linear relationship, as well as the significance (P < 0.05) of a quadratic term in the model. If a continuous variable did not have a linear relationship with the outcome of interest, it was subsequently categorized based on biologically relevant cut-points, or if appropriate, a quadratic term was retained in the model. Variables retained in the final model were assessed for collinearity through the examination of Pearson or Spearman rank-order correlation coefficients. When high correlation was found between variables (> 0.6), the most biologically appropriate variable was chosen for inclusion in the final model. Standardized residuals were generated and visually assessed for normality and homoscedasticity for the linear mixed regression model. If the residuals were heteroskedastic or not normally distributed, appropriate transformations were performed on the outcome of interest.

## Results and Discussion

Table 1 outlines the descriptive characteristics of trial animals. The CB group had a lower absolute number of animals in their second and third lactation, although this difference was not statistically significant. The twinning rate in each group was relatively low, with 2 twin calvings in the CB group and 1 in the NB group. Mean non-lactating days was numerically greater for NB cows, but this difference was not statistically significant. Days dry and its quadratic function were included in the final linear model however, as it was significantly associated with serum calcium (Table 2) and it had a confounding influence noted on the coefficients for treatment.

**Table 1.**
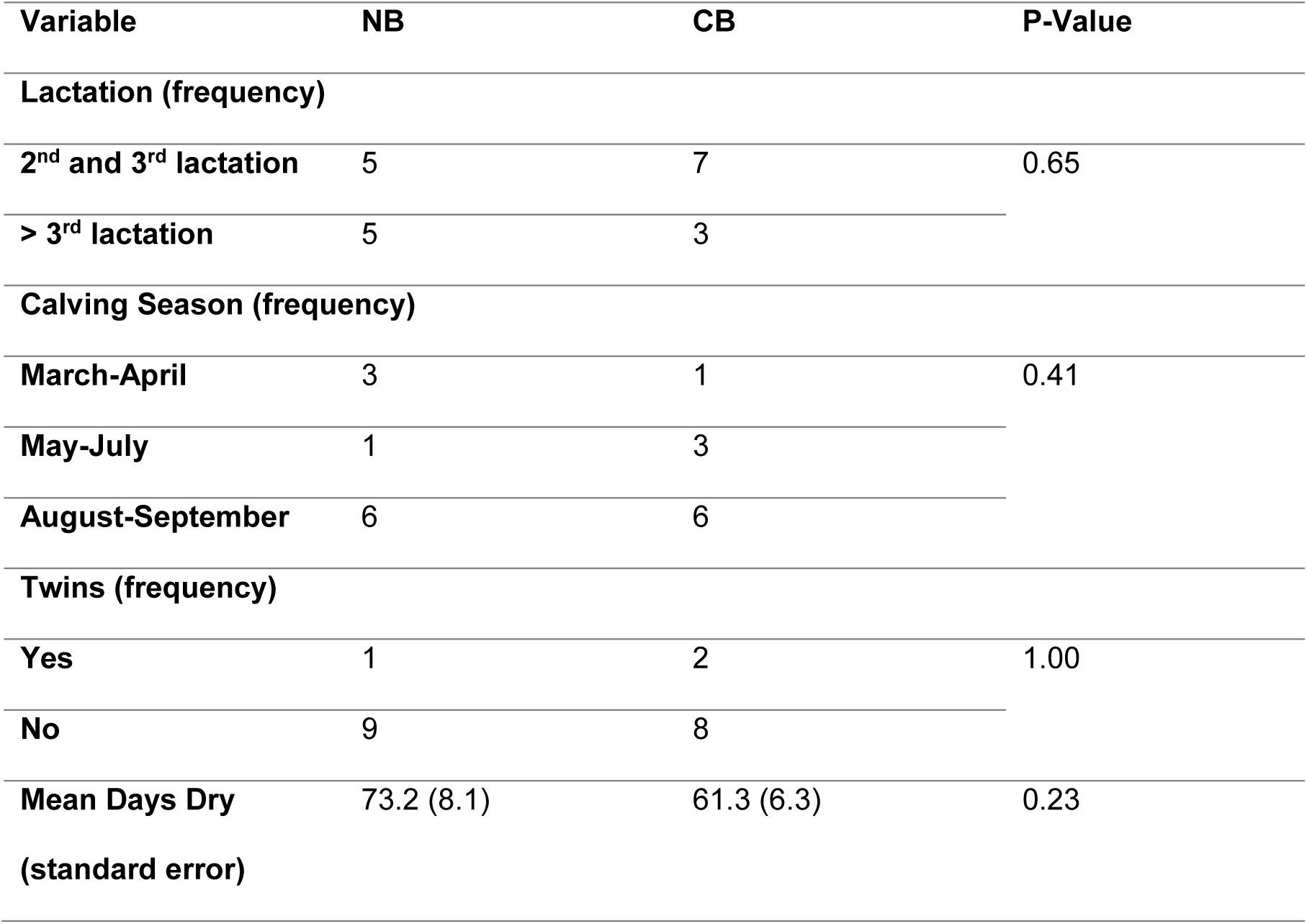
Descriptive demographic comparisons between a novel calcium bolus (NB) and a commercially available bolus (CB) of cows enrolled on a commercial dairy farm in southwestern Ontario, Canada.

**Table 2.** Multivariable, repeated measures linear regression model output evaluating the effect that the novel bolus (NB) relative to the commercially available bolus (CB) has on log-transformed serum calcium values within the first 24 hours following parturition.

|  | Coefficient | Standard Error | P-Value | 95% Confidence Interval |  |
| --- | --- | --- | --- | --- | --- |
| Treatment |  |  |  |  |  |
| CB | Referent |  |  |  |  |
| NB | 0.03 | 0.05 | 0.50 | -0.06 | 0.12 |
| Sample Time (h relative to parturition) |  |  |  |  |  |
| 0 | Referent |  |  |  |  |
| 1 | 0.18 | 0.05 | 0.001 | 0.08 | 0.29 |
| 6 | 0.07 | 0.06 | 0.23 | -0.04 | 0.18 |
| 12 | 0.04 | 0.06 | 0.52 | -0.08 | 0.15 |
| 13 | 0.01 | 0.06 | 0.80 | -0.10 | 0.13 |
| 24 | -0.01 | 0.06 | 0.82 | -0.13 | 0.10 |
| Calving Month |  |  |  |  |  |
| March-April | Referent |  |  |  |  |
| May-July | -0.04 | 0.07 | 0.54 | -0.17 | 0.09 |
| August-September | 0.16 | 0.05 | 0.002 | 0.06 | 0.27 |
| Days Dry | 0.01 | 0.00 | <0.001 | 0.01 | 0.02 |
| Days Dry (quadratic) | 0.00007 | 0.00002 | 0.001 | 0.00 | 0.00 |
| Cow Lactation: |  |  |  |  |  |
| 2<sup>nd</sup> and 3<sup>rd</sup> | Referent |  |  |  |  |
| **3<sup>rd+</sup>** | -0.08 | 0.05 | 0.11 | -0.17 | 0.02 |
| **Constant** | -0.07 | 0.15 | 0.65 | -0.36 | 0.23 |

Figure 1. a. outlines a descriptive comparison in serum calcium levels between NB and CB-treated animals. Of note, NB cows had lower serum calcium at parturition relative to CB animals, but the NB group had numerically higher serum calcium levels in subsequent sampling times (Figure 1.a.). Figures 1.b. and 1.c. outline raw mean (SD) serum phosphorus and magnesium levels measured in study cows, respectively. There were no significant differences between NB and CB groups for all blood metabolites (P > 0.05) measured on univariable analysis. Of note, there were 2 animals of 20 that were sampled slightly outside of the appropriate window (time ± 1 hour) as outlined in the study protocol for the 12-hour samples, both of which were in the NB treatment group. Analyses were conducted both including and excluding these animals, with no bearing on the outcome of the experiment. Therefore, the data for these animals remained in the analysis (N = 20) to avoid excessive model overfitting.

**Figure 1.**
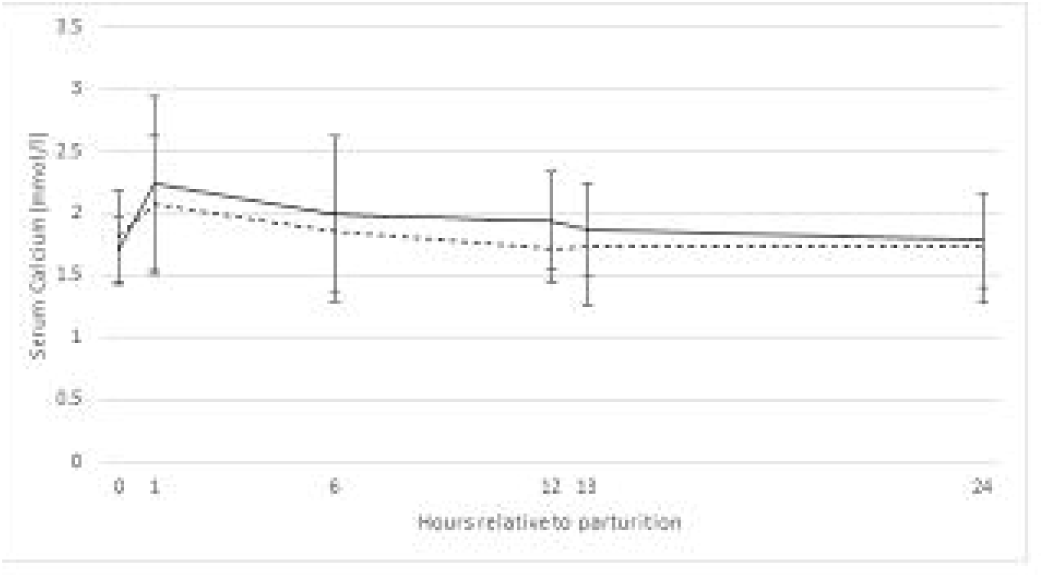
a. Raw mean (standard deviation) serum calcium (mmol/L) plot of novel calcium bolus (solid line) and commercially available bolus (dashed line) treatment groups over the first 24 hours following parturition.

**Figure 1.**
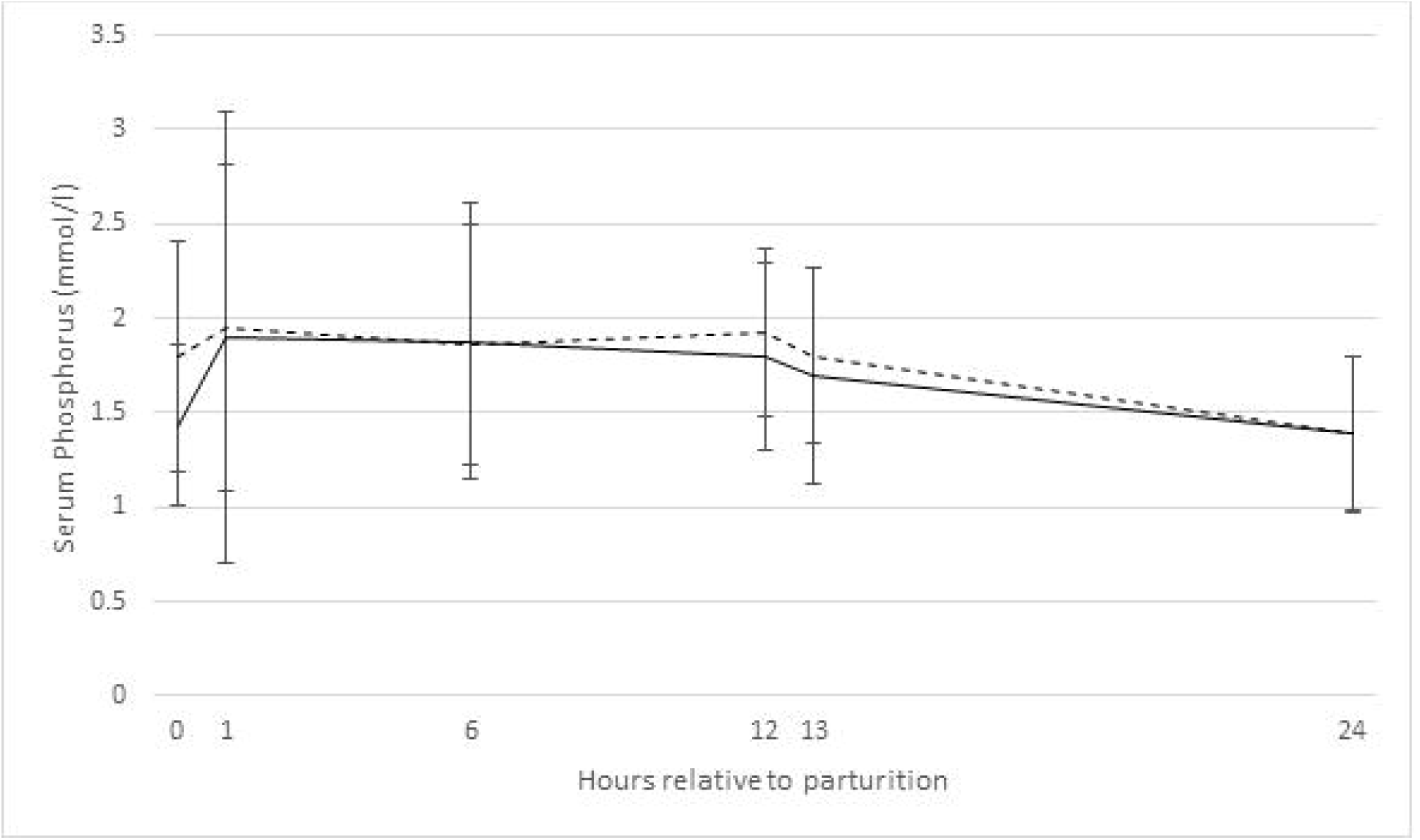
b. Raw mean (standard deviation) serum phorphorus (mmol/L) plot of novel calcium bolus (solid line) and commercially available bolus (dashed line) treatment groups over the first 24 hours following parturition.

**Figure 1.**
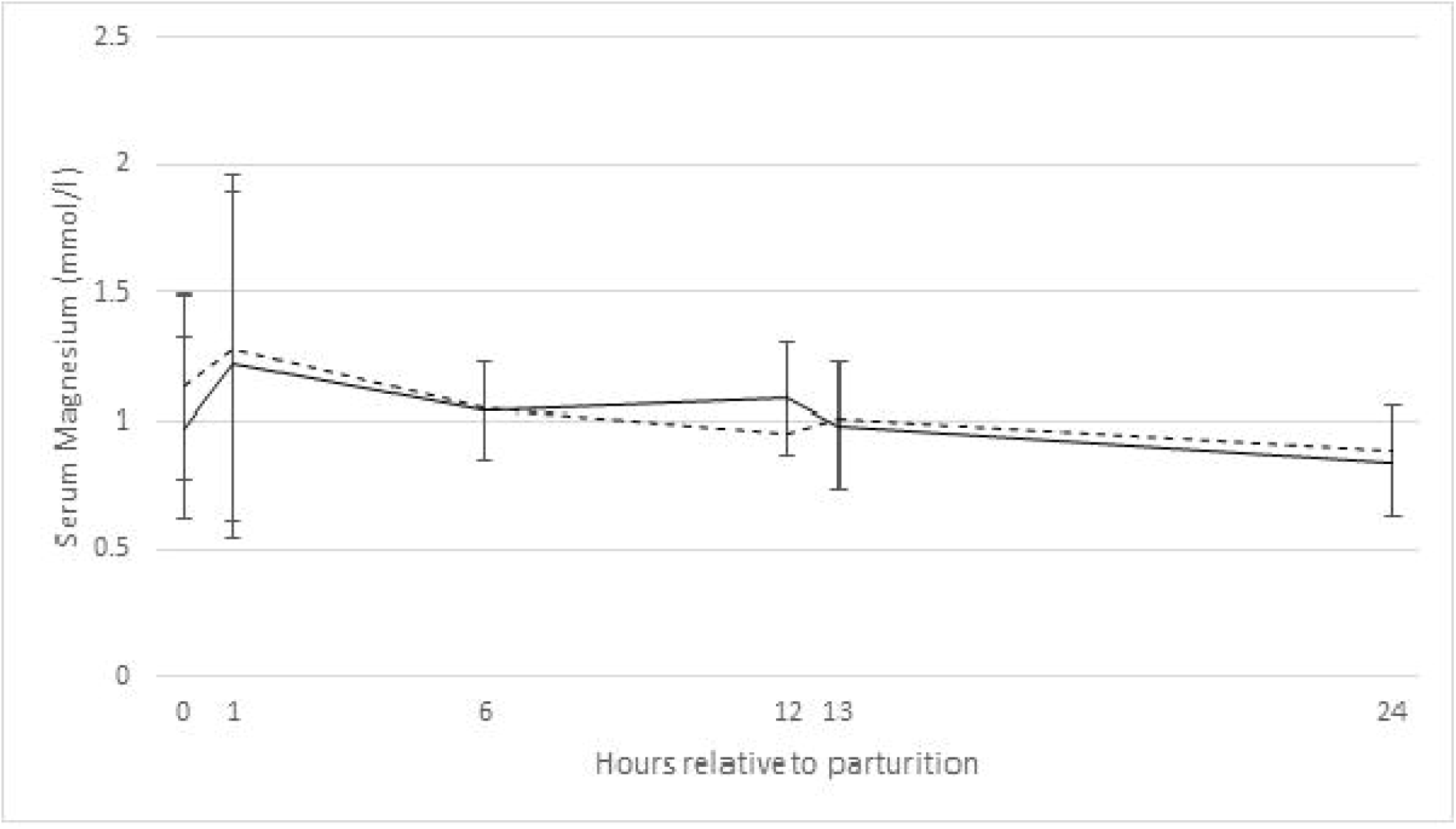
c. Raw mean (standard deviation) serum magnesium (mmol/L) plot of novel calcium bolus (solid line) and commercially available bolus (dashed line) treatment groups over the first 24 hours following parturition.

Residual analysis from raw serum calcium data showed considerable heteroskedasticity, therefore natural logarithms were calculated for each calcium value. Upon subsequent iterations of the model, residuals were homoscedastic and normally distributed with this transformation. Multivariable, repeated measures linear regression model output is given in Table 2.

Serum calcium levels did not differ at any point between CB- and NB-treated cows (P = 0.50). Serum calcium levels rose dramatically at the 1-hour mark following parturition; however, these levels subsequently dropped and were not significantly different from baseline (t = 0) levels for the remainder of the 24-hour sampling period. Cows that calved in the months of August and September had significantly higher serum calcium levels (P = 0.002) relative to cows calving in March and April. Older cows tended (P = 0.11) to have lower serum calcium levels. This variable was included in the model, as it had a significant confounding effect (> 20% change in coefficient value) on the treatment variable of interest (NB versus CB-treatment). Previous research has noted older cows to be at an increased risk of clinical milk fever ^1^. Overall, Figure 2 outlines serum calcium levels for CB- and NB-treated cows over the experimental period.

Calcium supplementation of cows around parturition has been studied extensively for the treatment and prevention of hypocalcemia^1,2,9,11–13^. Subclinical hypocalcemia (serum calcium levels < 2.0 mmol/l) is a pervasive issue in the modern dairy industry. All cows in the current study experienced subclinical hypocalcemia. Previous research has found that the incidence to be approximately 50% in mature (2^nd^ or greater lactation) Holstein cows ^4,14^. Indeed, the risk of displaced abomasum, ketosis, retained placenta, and metritis was 3.7, 5.5, 3.4, and 4.3 times more likely, respectively, in cows that experience subclinical hypocalcemia relative to normocalcemic cohorts ^15^. Overall, the costs associated with subclinical hypocalcemia are estimated to be approximately $125 per case ^2^. For the average 500 cow herd, assuming 65% of the herd is in their 2^nd^ lactation or greater, and a 50% incidence of subclinical mastitis, the cost would be approximately $20,312.50. Clearly, significant efforts should be made to prevent such losses.

Cows that received both calcium supplements experienced significant increases in serum calcium levels; however, these levels were not sustained for the remainder of the 24 hours following parturition. There has been extensive research into the provision of supplemental calcium to cattle around parturition^2,9,11–14^. Early studies by Goff and Horst^12^ found that cows supplemented with 128.8 g of calcium chloride as an oral drench experienced significant elevations in serum calcium up to 6 hours following treatment. Interestingly, the authors noted that stimulating the esophageal groove with vasopressin prior to calcium administration resulted in higher serum calcium concentrations than calcium chloride supplementation alone^12^. This rumen bypassing effect could improve overall calcium absorption and potentially explain the variable influence that solid boluses have on serum calcium concentration, despite having significantly higher calcium chloride levels relative to liquid products tested^16^.

Over and above the transient effects of supplemental calcium, the acidifying nature of the calcium chloride and calcium sulfate could influence systemic pH^12,17^, as the feeding of anionic diets to transition cows is a proven strategy to reduce the incidence of hypocalcemia^18^. These results are consistent with a recent study that found oral calcium supplementation effectively increased serum calcium levels at the 1 and 24 hours postpartum relative to untreated controls ^8,11^. One important difference between the current experiment and the cited studies was the use of negative control animals. As the current study was a pilot field-based trial, and calcium supplementation is a pervasive practice throughout the industry, it was difficult to recruit a suitable herd capable of an intensive sampling schedule that would allow cows at high risk for hypocalcemia to remain untreated. On the other hand, numerous studies evaluating postpartum, serial serum calcium levels in cows not receiving supplemental calcium either remain constant or significantly decrease relative to baseline levels at calving ^8,11,17,19,20^. Given the lack of a concurrent negative control, no definitive conclusions can be made.

The addition of vitamin D3 to the NB has the theoretical advantage of improving plasma Ca and P concentrations through intestinal absorption and bone resorption^21^. Though supplementation with exogenous vitamin D has significantly affected serum Vitamin D levels^21^, the effects on serum calcium levels have been variable^22–24^. The cited studies used higher amounts of vitamin D^22–24^ and included them in the diets of transition cows^23,24^. To this author’s knowledge, the effects of bolus supplementation of 50,000 IU vitamin D3 have not been studies, and the utility of inclusion in commercial preparations is unknown.

The manufacturing and marketing of novel supplements (i.e. mineral boluses, vitamins, botanicals) are not subjected to as rigorous a regulatory process relative to veterinary pharmaceutical products. As such, there is a paucity of peer-reviewed literature evaluating such novel products in the field. The present pilot study does display that the novel product is able to cause an elevation in serum calcium of recently calved cows of a magnitude like another commercially available, efficacious product of similar composition.

## Conclusion

Subclinical hypocalcemia is a prevalent disorder of the modern commercial dairy cow. The current pilot study found oral supplementation with two commercially available calcium boluses was associated with a significant increase in serum calcium levels at 1-hour post-calving. Producers and veterinarians wishing to affect a sustained serum calcium response in fresh cows should not rely on calcium boluses and instead focus on preventative management and nutritional practices.

## Acknowledgments

The authors would like to thank Dr. Ray Reynen (Attending veterinarian from Heartland Veterinary Services) and the participating farm and its workers whose hard work made this trial possible.

